# Ca^2+^ oscillations, waves, and networks in islets from human donors with and without type 2 diabetes

**DOI:** 10.1101/2021.12.08.471749

**Authors:** Marko Gosak, Richard Yan-Do, Haopeng Lin, Patrick E. MacDonald, Andraž Stožer

## Abstract

Pancreatic islets are highly interconnected structures that produce pulses of insulin and other hormones, maintaining normal homeostasis of glucose and other nutrients. Normal stimulus-secretion and intercellular coupling are essential to regulated secretory responses and these hallmarks are known to be altered in diabetes. In the present study, we used calcium imaging of isolated human islets to assess their collective cell behavior. The activity occurred in the form of calcium oscillations, was synchronized across different regions of islets through calcium waves, and was glucose-dependent: higher glucose enhanced the activity, elicited a greater proportion of global calcium waves, and led to denser and less fragmented functional networks. Hub regions were identified in stimulatory conditions, and they represented the most active islet regions. Moreover, calcium waves were found to be initiated in different subregions and the roles of initiators and hubs did not overlap. In type 2 diabetes, glucose-dependence was retained, but a reduced activity, locally restricted waves, and more segregated networks were detected compared with control islets. Interestingly, hub regions seemed to suffer the most by losing a disproportionately large fraction of connections. These changes affected islets from donors with diabetes in a heterogeneous manner.

## Introduction

A major part of pancreatic islets comprises collectives of nutrient-sensing and insulin-secreting beta cells (1, 2). They adapt to changes in metabolic demands by corresponding changes in intracellular stimulus-secretion coupling and despite their heterogeneity, under normal conditions they respond in a synchronized manner (3-5). This is facilitated by gap junctions that allow for the exchange of ions and small metabolites between beta cells and coordinate electrical activity across the islets in the form of propagating intercellular waves (6, 7). Noteworthy, gap-junctional communication is instrumental for appropriate glucose-induced insulin release, whereas its impairment is associated with disruptions in plasma insulin oscillations, similarly to models of metabolic diseases (8, 9). Therefore, intercellular coupling and its modulation are increasingly recognized as key to normal islet function (10-12), and potentially viable targets to improve insulin secretion (13, 14).

Changes in intracellular calcium concentration ([Ca^2+^]_i_) play a central role in intracellular stimulus-secretion coupling and most of our knowledge on [Ca^2+^]_i_ signaling in islets derives from mouse models. Although mouse and human islets share key structural and functional features, the intracellular electrical and [Ca^2+^]_i_ activity in human beta cells are more variable and differ from those of mouse islets (2, 12). Furthermore, the collective beta cell rhythmicity is much less explored in human islets. Although there is a similar degree of gap junctional coupling in mouse and human beta cells (15), the unique cytoarchitecture of human islets and less coherent [Ca^2+^]_i_ activity may explain why distinguished [Ca^2+^]_i_ oscillations and waves have been more difficult to detect than in mouse islets (16-20). Advances in optogenetics and computational tools have allowed assessment of multicellular beta cell behavior in mice, revealing that the mediating [Ca^2+^]_i_ waves are initiated from specific subregions of the islet that are relatively stable in time. These subpopulations are also called pacemaker or leader regions and are defined by local excitability and metabolic profiles (21-25). In human islets, the synchronization patterns are similar but more clustered than in mice and the collective dynamics was found to change in disease (20, 24, 26).

Our increasing awareness of the importance of intercellular communication has stimulated the development of advanced computational approaches. Network science has provided powerful tools to assess collective cellular behavior in islets (20, 22, 24, 27-29). Typically, functional beta cell networks are constructed from thresholded pairwise correlations of [Ca^2+^]_i_ signals and the subsequent analyses show that the beta cell collectives organize into efficient, modular, and heterogeneous networks (22, 27, 30), in which a subset of metabolically highly active hub cells are believed to crucially enhance the communication capacities of intercellular networks (22, 31). The complex hub-like connectivity patterns arise from islet-wide [Ca^2+^]_i_ dynamics and are influenced by cellular heterogeneity, heterogeneous intercellular interactions, and interactions of cells with the environment (32). However, it is not entirely clear to what extent specialized subpopulations of cells contribute to synchronized network dynamics, persistency, and initiation of intercellular signals, and how these parameters change in type 2 diabetes. Particularly the reports about pacemaker cells and their relation to hub cells are inconsistent and require further clarification (33, 34).

Here we combined camera-based [Ca^2+^]_i_ imaging of human islets from organ donors with and without type 2 diabetes with classical physiological and advanced network analysis to investigate whether and how glucose controls [Ca^2+^]_i_ oscillations and waves, how the [Ca^2+^]_i_ dynamics relates to functional connectivity in islets, and how these parameters change in diabetes.

## Materials and Methods

### Islet isolation

Human islets were isolated in the Alberta Diabetes Institute IsletCore (35) or the Clinical Islet Laboratory (36) at the University of Alberta, with appropriate ethical approval from the University of Alberta Human Research Ethics Board (Pro00013094; Pro 00001754). Islets were hand-picked and cultured in the RPMI-1640 medium (Life Technologies) containing 7.5 mmol/L glucose, 10 % fetal bovine serum, and 5 % penicillin/streptomycin overnight or DMEM (Gibco), containing 5.5mmol/L glucose, 10 % fetal bovine serum, and 5 % penicillin/streptomycin. Donor characteristics and islet secretion data are presented in Supplementary table 1.

### Calcium imaging

Intact human islets were incubated in culture medium containing 5μM Fluo-4 (Invitrogen) for 60 min. Islets were mounted into a custom recording chamber and perifused with RPMI-1640 growth medium supplemented with 10 % fetal bovine serum and 5 % penicillin/streptomycin and glucose as indicated. The cells were continuously perifused with extracellular solution at a bath temperature of ∼32 °C. Imaging was performed on a Zeiss SteREO Discovery V20 upright microscope with ZEN acquisition software. Fluorescence was excited at 488 nm at 10 % light intensity using an LED light source. Images were captured with an AxioCam MRm CCD camera at 0.33 Hz at 200 ms exposure and recorded for 1 hour. 12-bit 1388 × 1040 pixels images were captured where each pixel was 1.02 um x 1.02 um.

### Image analysis and time series processing

The [Ca^2+^]_i_ dynamics in islets was analyzed off-line employing ImageJ (37) and custom-made scripts in Python. Small movements of islets in the chamber were removed with a stabilization algorithm and a 3D Gaussian filter was applied to denoise the recording. The *F*/*F*_0_ ratio of the fluorescence (*F*) with respect to initial fluorescence (*F*_0_) was calculated. The image was then binarized and subdivided into a square mesh, i.e., islet sub-regions (ISR), with an edge size of 15 µm, since the discrimination of individual cells was not possible. The mean greyscale values from all frames were exported from each ISR and the extracted [Ca^2+^]_i_ traces were processed with a high-pass filter to retrieve baseline trends and possible artefacts. This enabled a binarization of [Ca^2+^]_i_ signals, which were then used to calculate the signaling parameters, and for the extraction of [Ca^2+^]_i_ waves. The latter was achieved by means of a space-time cluster analysis, as described previously (38). We calculated the average wave sizes and identified the initiating ISRs. All ISRs wave were ranked in accordance with their order of activation in a [Ca^2+^]_i_ wave. The ranks were rescaled to unit interval, with the lowest initiation rank signifying the first responding ISR. The average initiation rank of a given ISR was calculated as the average over all Ca^2+^ waves. How [Ca^2+^]_i_ signals were processed is shown in Supplementary figure 1 and how the [Ca^2+^]_i_ waves were extracted is presented in Supplementary video S1.

### Functional network analysis

Filtered [Ca^2+^]_i_ signals were used for the correlation analysis and the subsequent network construction. Two ISRs were functionally connected if the pairwise correlation coefficient between the *i*-th and *j*-th ISR, *R*_*ij*_, exceeded a given threshold value, as described elsewhere (20, 27, 29). We used *R*_*ij*_>0.7 for the establishment of networks and diagnosed them with conventional network metrics. We calculated the degree of individual ISRs, the average degree, and the relative degree distribution. To quantify network integration, we calculated the relative largest component, which signifies the fraction of cells that are either directly or indirectly connected and how many of them are isolated. To quantify functional segregation of islets, we calculated modularity, which measures how effectively the network can be partitioned into submodules. Higher values of modularity indicate more segregated network structures (29).

### Statistical analysis

Differences in parameters between 8 mM and 12 mM glucose were assessed using the t-test or the Mann-Whitney test. Two-tailed ANOVA was used to compare the parameter values between islets from control and normal donors in 8 mM and 12 mM glucose. One-way ANOVA or the Kruskal-Wallis test were used to compare active time, initiation rank, and node degree for ISRs with the upper, intermediate, and lower third parameter values. Statistical significance is indicated with asterisks, as follows: * p<0,05; ** p<0,01; *** p<0,001. The exact values are indicated for p-values between 0,05 and 0,1.

## Results

We analyzed [Ca^2+^]_i_ signals in isolated human islets from donors with and without type 2 diabetes after stimulation with glucose. Our protocol consisted of an initial ∼15 min of sub-stimulatory 3 mM glucose, a subsequent stepwise increase to stimulatory 8 or 12 mM glucose for ∼45 min, and a decrease back to 3 mM glucose. We analyzed altogether 40 islets from 6 normal donors and 36 islets from 6 donors with diabetes (Supplementary Table 1). For the analysis of various signaling parameters, we selected 20-30 min intervals of stable [Ca^2+^]_i_ activity from each islet. Since the spatial resolution of the recording setup did not enable identification of individual cells, the islets were subdivided into islet sub-regions, ISRs, as shown in Figures 1A and 1E.

**Figure 1.**
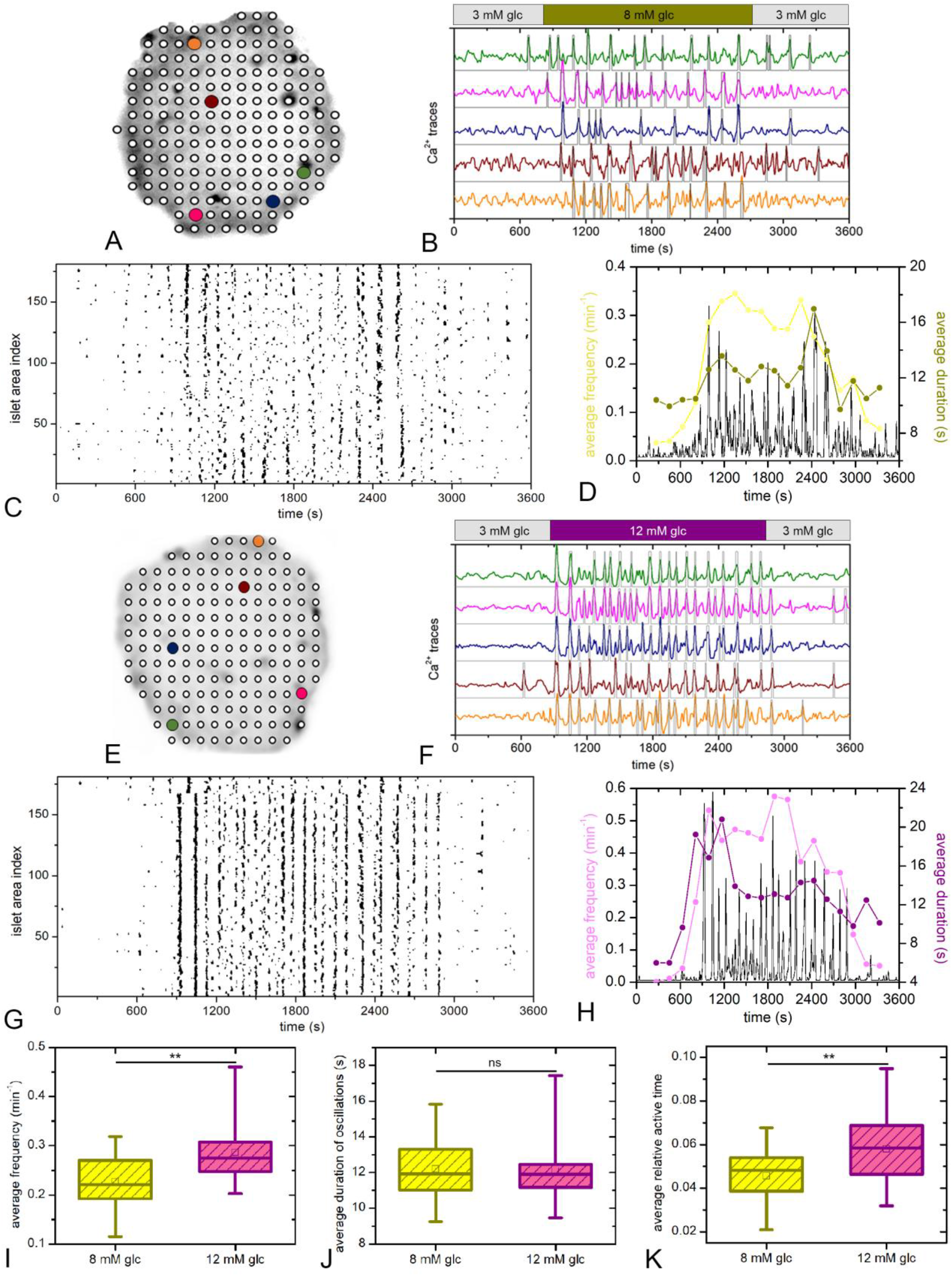
Glucose-stimulated [Ca^2+^]_i_ oscillations in human islets. Islets segmented into ISRs (A and E) and their [Ca^2+^]_i_ activity in five representative areas, as indicated by the colors of dots and traces (B and F). Grey lines denote binarized signals corresponding to specific traces. Binarized activity of all ISRs as a function of time (C and G). The evolution of the average frequency and duration of oscillations in a sliding temporal window (width 270 s, step 180 s) for the selected islets (D and H). The grey line indicates the mean-field signal of binarized traces. Panels (A-D) and (E-H) refer to 8 mM and 12 mM glucose stimulation, respectively. Panels (I-K) show [Ca^2+^]_i_ oscillation characteristics for both glucose concentrations, pooled from all islets (20 islets and 2697 regions in 8 mM glucose, 20 islets and 3205 regions in 12 mM glucose). In panels (I, J, K) boxes determinate the interval between the 25th and the 75th percentile, whiskers denote the 10th and the 90th percentile, lines within the boxes indicate the median, and small squares stand for the average value.

### Glucose stimulates [Ca^2+^]_i_ oscillations in human islets

The processed [Ca^2+^]_i_ activities of selected regions when glucose increased from 3 mM to 8 mM or 12 mM for ∼30 min are shown in Figures 2B and 2F, respectively. To provide a more quantitative and detailed insight into the [Ca^2+^]_i_ activity, we show raster plots of binarized activity for all ISRs within an islet (Figures 2C and 2G), along with the temporal evolution of the average oscillation frequency and duration (Figures 2D and 2H). Both stimulatory glucose levels led to a relatively stable plateau phase with sustained [Ca^2+^]_i_ oscillations. Pooled data are shown in Figures 1I-K. An approximately 20 % higher active time was observed in 12 mM compared with 8 mM glucose, solely due to a higher average frequency. The average duration of oscillations under both stimulation levels was ∼12 s, whereas the average frequencies were 0.22 and 0.27 min^-1^ for 8 mM and 12 mM, respectively (Figures 1 I and J). Moreover, we characterized ISR activations after switching from 3 mM glucose to stimulatory levels. We considered an ISR as activated when the first [Ca^2+^]_i_ oscillation occurred after the rise in glucose. Higher glucose increased the rate of recruitment by ∼30 % (Supplementary figure S2).

**Figure 2.**
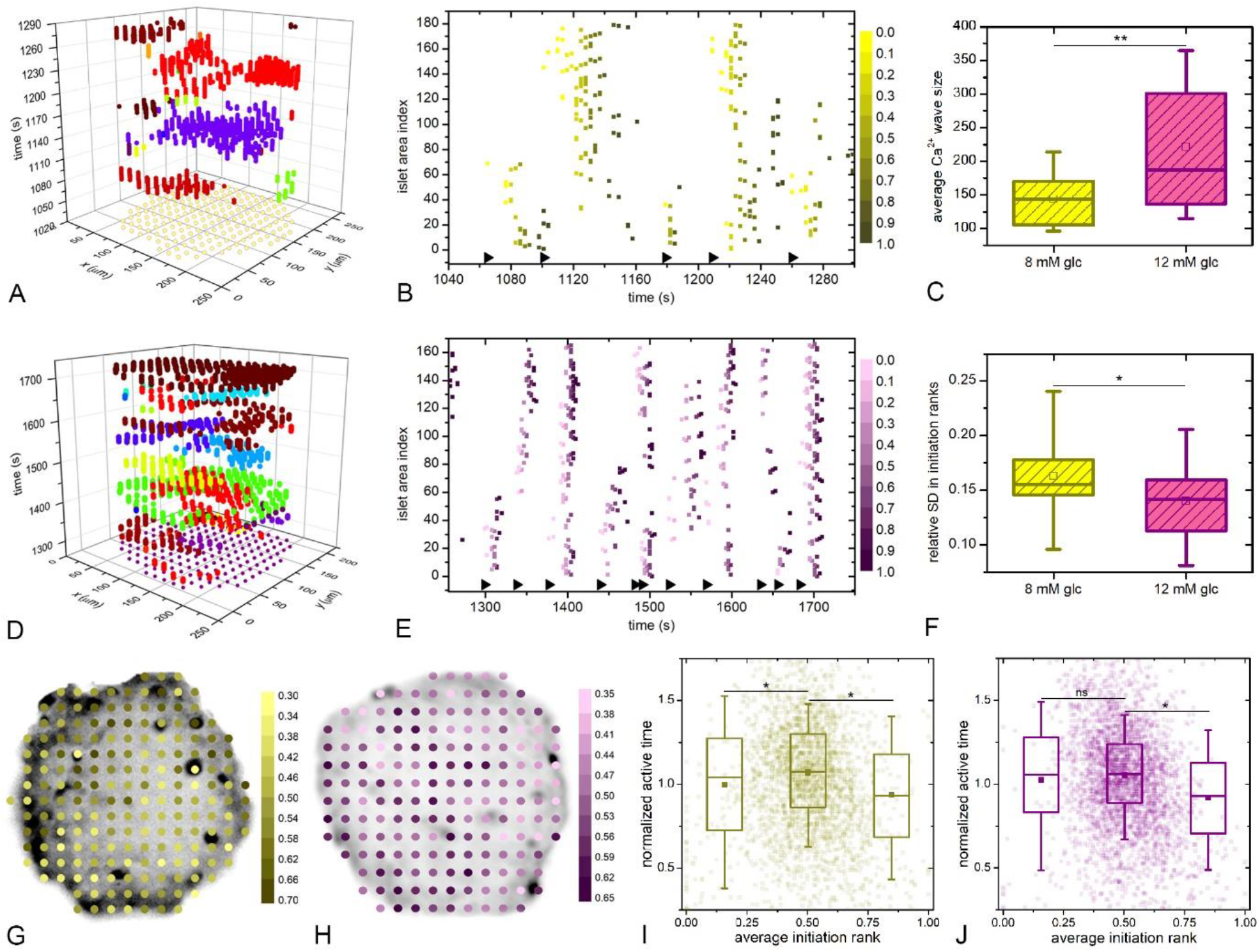
Characteristics of [Ca^2+^]_i_ waves in human islets. Space-time [Ca^2+^]_i_ clusters for given time intervals in 8 mM (A) and 12 mM glucose (D). Individual [Ca^2+^]_i_ waves are denoted by different colors, the dots in the *x*-*y* plane denote the spatial coordinates of individual ISRs within an islet and time is indicated in the *z*-axis. The corresponding raster plots of oscillation onsets for the waves shown in A and D, with normalized color-coded activation ranks for a given oscillation in 8 mM (B) and 12 mM glucose (E). The black triangles in the bottom indicate the beginnings of individual waves. Panels (C) and (F) show the comparison between average wave sizes and relative SDs in initiation ranks in 8 mM vs. 12 mM glucose, pooled from all islets in the given group. Box charts are defined as in Fig. 1. Panels (G) and (H) show ISRs with color-coded values of average initiation ranks for 8 mM and 12 mM glucose, respectively. The average normalized active time as a function of average initiation ranks in 8 mM (I) and 12 mM (J) glucose. Semi-transparent dots signify values of all ISRs from different islets and the box-plots signify the lowest, the intermediate, and the highest one third of initiation rank parameter values.

### Glucose modulates the organization of [Ca^2+^]_i_ waves in human islets

Plotting [Ca^2+^]_i_ oscillations as individual events in space-time demonstrated that [Ca^2+^]_i_ waves are the mechanism of interregional synchronization (Figures 2A and 2D). We then calculated the average sizes of [Ca^2+^]_i_ waves and the relative SD of initiation ranks of ISRs for all islets and for whole intervals of sustained activity. The latter was used as a measure for the persistency of wave courses and initiator regions. Very small [Ca^2+^]_i_ waves were excluded from further analysis, since many small ISR activations were due to unavoidable noise in the [Ca^2+^]_i_ signals. Within the remaining [Ca^2+^]_i_ waves, we ranked all ISRs by the sequence of activations, as illustrated with raster plots in Figures 2B and 2E. Under 8 mM glucose, [Ca^2+^]_i_ waves of very different sizes were observed, whereas under 12 mM glucose, the waves were less heterogeneous and more often encompassed substantial parts of the islet. Moreover, [Ca^2+^]_i_ wave initiation occurred in different ISRs and their courses did not follow the same sequence of activation. The initiator role often changed from one event to another. Comparing the relative SD of initiation ranks indicated that the sequence of activations was more persistent and thus the initiator regions more stable in 12mM glucose.

In Figures 2G and 2H, ISRs are presented with color-coded values of average initiation ranks. It can be observed that for both glucose concentrations the wave initiating regions are scattered across the islet without an obvious pattern (Supplementary video S1). Moreover, in Figures 2I and 2J the average active times are plotted as a function of the relative initiation ranks. Only the values of ISRs that participated in at least 10 % of all [Ca^2+^]_i_ waves were considered, and all values were normalized with the corresponding average values of active times to enable pooling of data from different islets. First, it can be noticed that the relative initiation ranks are scattered between 0.25 and 0.75, whereas ISRs with very low or high initiation ranks were detected only exceptionally. This suggests that the roles of ISRs in the wave initiation process were rather dynamic, with a given ISR sometimes acting as an initiator and sometimes being activated among the last within a wave. Analyzing the relationship between an ISR’s active time and its role in wave initiation revealed that the waves were rarely triggered by ISRs with the lowest active times.

### Glucose controls the functional connectivity networks in human islets

To further characterize the collective activity, we constructed functional connectivity maps for both glucose concentrations. [Ca^2+^]_i_ signals from all ISRs were statistically compared in a pair-wise manner to build correlation matrices (Figures 3A-D), which were then thresholded to obtain functional networks. For both stimulation levels, the network architectures appeared rather regular and lattice-like (Figures 3C and 3F), in contrast to β-cell networks inferred from mouse islets which exhibit more long-range connections (22, 24, 27-29). Comparing networks in 8 mM and 12 mM glucose showed that under higher stimulation, the networks were denser (higher average degree, Figure 3G), more cohesive (higher relative largest components, Figure 3H), and less fragmented (lower modularity, Figure 3I). These properties reflect the fact that the [Ca^2+^]_i_ waves provoked by 12 mM glucose typically encompassed larger parts of an islet, whereas in 8 mM glucose, they were typically limited to smaller parts of an islet. The node degree distributions in Figure 3J show that under both glucose concentrations, the islet networks were rather heterogeneous. A relatively small fraction of ISRs existed that were very well connected, with so-called hub areas being functionally correlated to up to 40 % of all other ISRs.

**Figure 3.**
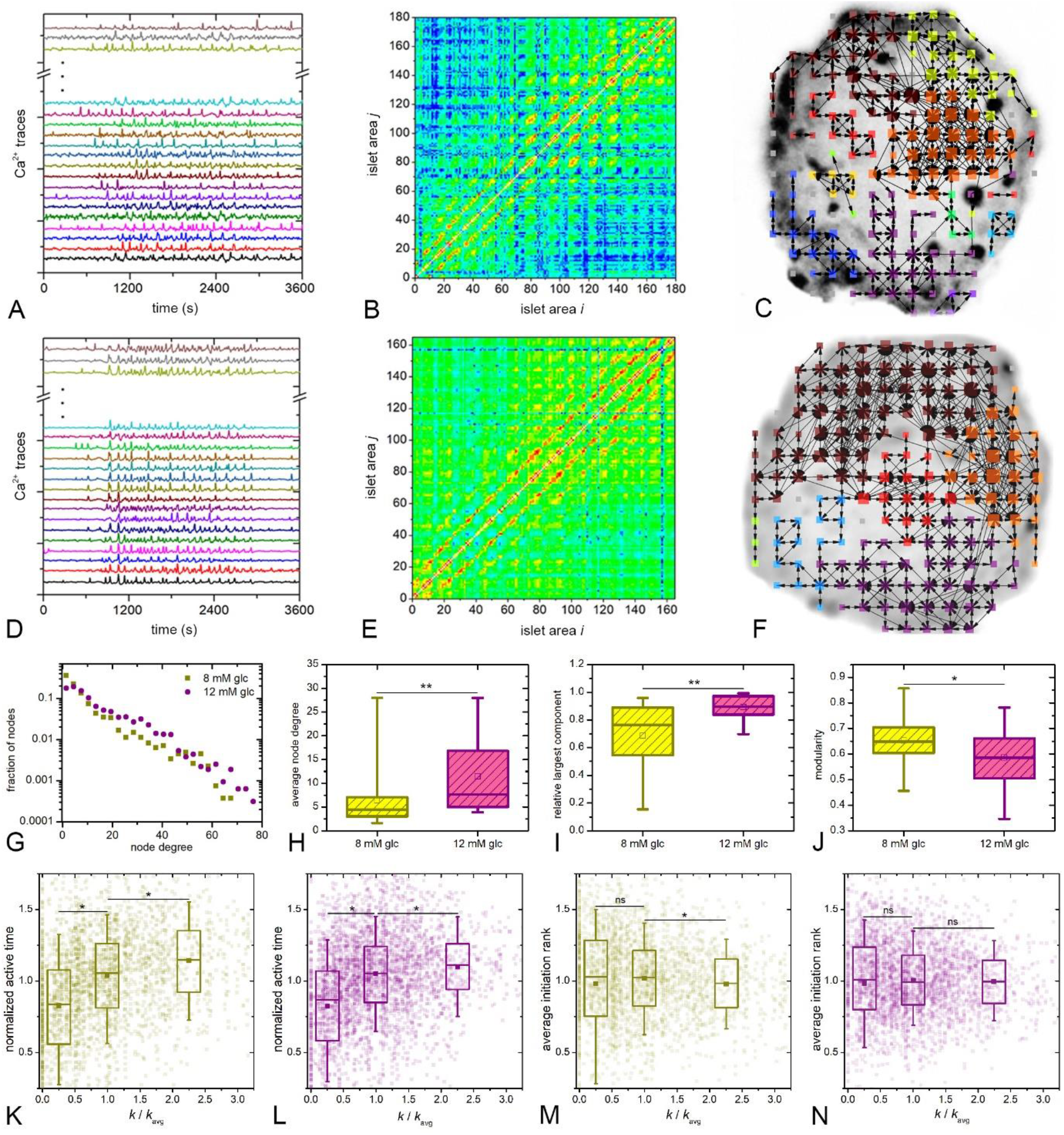
Functional connectivity networks in human islets. Mapping the functional connectivity of human islets: from the recorded [Ca^2+^]_i_ dynamics of all ISRs (A, D) to pair-wise correlation matrices (B, E) and networks (C, F). Panels (A-C) and (D-F) refer to stimulation with 8 mM and 12 mM glucose, respectively. G) Node degree distributions combined for all islets stimulated by a given glucose concentration. Panels (G-J) feature the combined results of the network metrics for all islets stimulated by a given glucose concentration: degree distribution (G), average network degree (H), relative largest component (I), and modularity (J). In panels (K-N) the relationship between relative node degree and either relative active time (K,L) or average initiation rank (M,N) are shown. Transparent dots indicate individual ISRs, whereas the box-plots denote the least connected cells (*k*/*k*_avg_≤0.5), intermediately connected cells (0.5< *k*/*k*_avg_ ≤1.5), and the most connected cells (*k*/*k*_avg_ >1.5), Box charts are defined as in Fig. 2. The results were obtained on the basis of 2697 and 3205 ISRs for 8 mM and 12 mM glucose, respectively.

To systematically compare ISRs with different node degree values with respect to their active times and wave initiation ranks, we pooled data from altogether 2697 regions for 8 mM glucose and 3205 regions for 12 mM glucose and divided them into three groups based on node degrees. In Figures 3K-3N the relationships between the above parameters are shown for all ISRs with semi-transparent dots, whereas the three box-plots indicate the least connected, the intermediately connected, and the most connected ISRs. Note that the node degrees and active times were normalized with the average values of the given islet to enable the pooling of ISRs from islets of different sizes and average activities. Evidently, for both glucose concentrations, the least active ISRs had the lowest number of connections, whilst the most active cells harbored the most connections (Figures 3K and 3L). The differences between active times between the least and most connected ISRs in 8 mM and 12 mM glucose were around 40 % and 30 %, respectively. In contrast, no clear relationship between the average initiation rank and node degree could be found (Figures 3M and 3N). The values of initiation ranks differed by less than 2.5 % from the average, irrespective of node degree and glucose level. This corroborates the findings in Figure 2 that the initiation of [Ca^2+^]_i_ waves was dynamic and occurred in different parts of an islet, in this case by either weakly or highly functionally connected ISRs with roughly the same probability.

### Changes in [Ca^2+^]_i_ activity in islets from patients with type 2 diabetes

Figure 4A-F shows the activity of two islets from the same donor with type 2 diabetes. While the [Ca^2+^]_i_ responses in one islet are similar to those in normal islets, in the other there is less activity, the [Ca^2+^]_i_ waves are smaller, and the functional networks are sparser and much more segregated. To summarize the general behavior of islets from donors with diabetes, Figures 4G-N show data for all islets examined. To facilitate comparison with control islets, boxplots showing the values for normal islets are also displayed in a thinner and brighter form. Notably, for most parameters, the islets from donors with type 2 diabetes exhibited a similar glucose-dependency as normal islets. However, in both glucose concentrations they were on average significantly less active (by 30-35 %), predominantly due to a lower frequency of oscillations (Figures 4G-I). Interestingly, the decrease in average active time could be attributed in part to a much higher proportion of ISRs with a very weak activity. More specifically, while the islets from control donors exhibited a normal-like distribution of ISR activity, in islets from donors with diabetes, the distribution was right-skewed and exhibited a higher fraction of weakly active ISRs (Figures 4O-Q), implying that the functioning of some regions was much more impaired than of others. The islets from donors with diabetes also responded to stimulatory glucose levels more slowly, especially to 8 mM (Supplementary Figure S2). Moreover, their [Ca^2+^]_i_ waves were smaller and more localized compared to normal responses in both glucose concentrations (Figure 4J) and the wave initiation sites and their courses were more erratic, as the relative SD in initiation ranks was significantly higher (Figure 4K).

**Figure 4.**
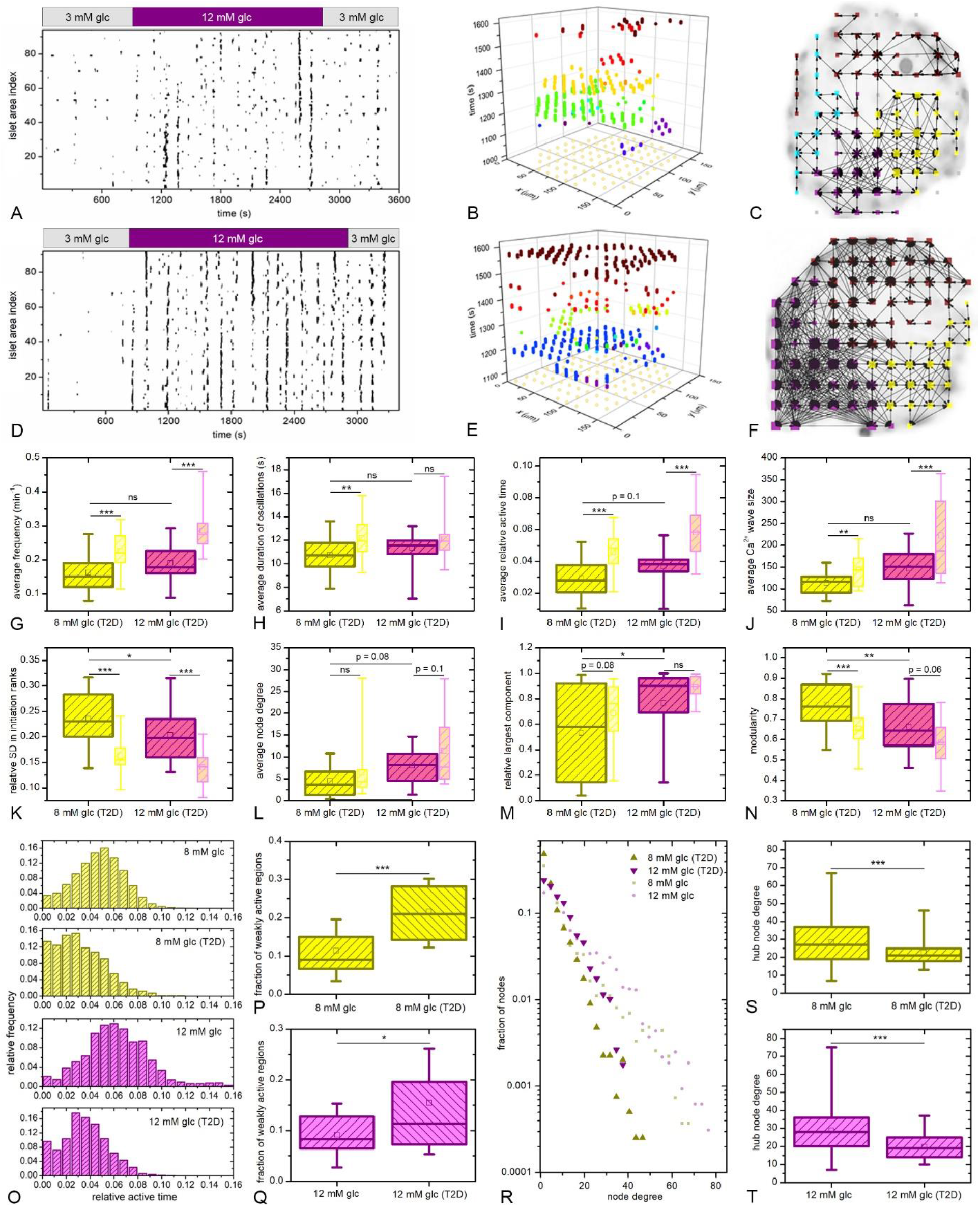
Changes in [Ca^2+^]_i_ activity in type 2 diabetes. Binarized activity of all ISRs for two different islets from the same donor with type 2 diabetes (A, D). [Ca^2+^]_i_ waves presented as space-time plots for selected intervals of the same two islets (B, E) and the resulting functional islet networks (C, F). Box-plots in panels G-N show different [Ca^2+^]_i_ signaling and islet network characteristics for both glucose concentrations pooled from all islets from donors with type 2 diabetes. The thinner and brighter plots show the data from non-diabetic islets from Figures 1-3 for comparison. The distribution of relative active times for both glucose concentrations, separately for islets from normal donors and donors with type 2 diabetes. The plots show the pooled data for all islets in a given group. Panels P and C represent the relative fraction of weakly active ISRs in all four groups. An ISRs was considered as weakly active if the active time was below 33 % of the average active time in the given islet. Node degree distributions combined for all islets under a given glucose concentration (R). Bright small dots show data from non-diabetic islets. Panels S and T show the node degrees of hub ISRs that were connected with at least 10 % of the islet, separately for islets from non-diabetic donors and donors with diabetes, for both glucose concentrations. The pooled data encompasses 23 islets (3971 ISRs) and 13 islets (2256 ISRs) from diabetic donors for 8 mM and 12 mM glucose stimulation, respectively. All box-plot charts are defined as in Figures 1-3.

The observed differences in the spatiotemporal [Ca^2+^]_i_ activity were accompanied by changes in functional connectivity. The degree distributions shown in Figure 4R indicate that islets from donors with diabetes lack very well-connected ISRs, i.e., hubs. Due to a fewer number of large [Ca^2+^]_i_ waves, there was no long-range coordination of intercellular signals, and consequently hub ISRs were less pronounced. In both glucose concentrations the average degree of highly connected ISRs, i.e., regions which were connected to at least 10 % of all ISRs in the given islet, was significantly lower in islets from donors with type 2 diabetes (Figures 4S and T), while the average node degree over all ISRs was not significantly lower (Figure 4L). Since the functional networks in human islets exhibited a rather lattice-like structure, the average number of connections per ISR reflects essentially the local synchronization level, which was not altered in diabetes. Moreover, no significant differences were found between the relative largest components of islets from donors with and without diabetes, but there was a much higher fraction of islets with a very low level of cohesion in both glucose concentrations in the islets from donors with diabetes (Figure 4M). Resulting from of a lack of global [Ca^2+^]_i_ waves, the islets from diabetic donors were also functionally more segregated, although the difference seemed to be more pronounced under 8 mM glucose (Figure 4N). Finally, the relationships between active time, initiation ranks, and number of functional connections in islets from donors with diabetes were very similar to what was observed in islets from non-diabetic donors (Supplementary figure S3).

## Discussion

In the present study we aimed to advance our understanding on beta cell spatiotemporal [Ca^2+^]_i_ patterns in human islets by systematically quantifying the glucose-dependent collective activity through the utilization of intensive computational approaches. As recently reviewed elsewhere (12), previous studies involving [Ca^2+^]_i_ imaging in human islets have reported globally (17, 18) or locally (19) synchronized [Ca^2+^]_i_ oscillations, the presence of some [Ca^2+^]_i_ oscillations without quantifying them or their synchronicity in detail (15, 20, 22, 39), or no clearly discernible oscillations (16). Two studies have evaluated [Ca^2+^]_i_ oscillations in human islets quantitatively, one by using a single stimulatory glucose concentration (11 mM) (19) and the second by using two different concentrations (11 mM and 16.7 mM) (18). The frequencies of oscillations in our study are comparable to these previous studies and are in the range of 0.1 to 1 min^-1^. Comparing the difference between 8 mM and 12 mM glucose in our study and the difference between 11 mM and 16.7 mM glucose in the study by Martin et al. (18), it seems that human islets respond to increasing glucose by first increasing the frequency and then the duration of oscillations, which qualitatively resembles the behavior in mouse islets (30). However, further studies are required to clarify the exact nature of [Ca^2+^]_i_ oscillations in human islets and the fact that the active time in human islets seems to be almost an order of magnitude lower than in mice (30).

To our knowledge, no previous study quantified the characteristics of [Ca^2+^]_i_ waves and functional networks to the extent that we could make a comparison with our findings. The observed behavior of [Ca^2+^]_i_ waves and functional networks, as well as their response to increasing glucose, can be attributed to a relatively less pronounced cellular heterogeneity at elevated glucose, which in turn diminished the spatiotemporal variations in excitability, making the courses of [Ca^2+^]_i_ waves more stable. Moreover, the distribution of connections between ISRs was found to obey an exponential function, indicating a lower heterogeneity of human islet functional networks compared to mouse islets, in which the degree distribution typically follows a truncated power-law (22, 27, 30). This discrepancy can at least partly be attributed to structural and functional differences between mouse and human islets and a somewhat different nature of [Ca^2+^]_i_ wave organization (1, 12, 16, 40). More specifically [Ca^2+^]_i_ waves in mouse islets commonly spread over the whole islet (6, 25), whereas in human islets we found them to be much more heterogeneous in size and often encompass only smaller regions (Figure 3). Together with donor and method heterogeneity (41, 42), this may also help explain why previous studies were often unable to detect [Ca^2+^]_i_ oscillations and waves in human islets and why the existing estimates of their characteristics differ (12).

We gave particular emphasis to the identification of specialized functional subpopulations of cells and their characteristics in terms of [Ca^2+^]_i_ activity. The heterogeneous distributions of functional connections among cells support the existence of so-called hub regions. Although the extent to which they coordinate the collective response and the temporal persistence of their role are not yet clear (33, 43), our finding that hub ISRs have the highest active times is consistent with recent findings in mouse islets (30) and the view that hub cells are metabolically highly active and exert a greater control over the electrical response of neighboring cells (4, 22, 23, 31). Furthermore, previous studies in mouse islets have shown that [Ca^2+^]_i_ waves originate from pacemaker regions with elevated excitability (6, 25), whereby the pacemaker cells were identified as metabolically less active with higher intrinsic frequencies (23). In our analyses we did not identify a clear relation with the activity of ISRs, except that the regions with low activity seldom initiated the waves. The triggering of waves was rather changeable and switched between areas scattered throughout the islets, especially under lower glucose, where intercellular variability was more pronounced. These concepts parallel with the idea that presence of pacemakers in networks of excitable cells reflects emergent dynamical behavior (44). Most importantly, we noticed no clear relationship between wave-initiation and hub ISRs, i.e., the waves could be triggered from either weakly or highly functionally connected ISRs. Apparently, hub and pacemaker cells are both genuine features of human islets, but their roles do not necessarily overlap and should not be considered equal. Importantly, these two populations also seem to differ from so-called first responders, i.e., cells that drive the first transitory phase of [Ca^2+^]_i_ elevation (30, 45).

Lastly, we investigated [Ca^2+^]_i_ signals in islets from donors with type 2 diabetes. To our knowledge, no detailed studies on how the [Ca^2+^]_i_ oscillations, waves, and coordinated network activity change in diabetes in humans have been conducted, yet a body of circumstantial evidence suggests that impaired coupling plays an important role (11, 12) and that a decline in coordinated islet activity is observed in diabetic conditions (20, 24, 26). Our analyses revealed that in these islets the active time is indeed decreased, mostly due to decreases in frequency of oscillations and heterogeneous dysfunction of some regions within the islets, the [Ca^2+^]_i_ waves are smaller, and the islet functional networks more segregated compared to normal islets. Notably, in our hands the loss of hub regions was accompanied by a strong decrease in [Ca^2+^]_i_ activity and a lack of global [Ca^2+^]_i_ waves, which supports the idea that the presence of hub regions may be necessary for normal intercellular synchronization of [Ca^2+^]_i_ activity (15, 20, 26). Finally, it should be noted that we observed a rather large variability in responses of islets from both normal and diabetic donors, with some of the latter exhibiting striking pathophysiological changes and other behaving rather normally. This supports the view that heterogeneity is an important aspect of normal islet functioning (4) and influences the susceptibility of different islets towards diabetogenic insults (46). Therefore, it should be considered as an important aspect when conducting research on human islets in general, also far beyond [Ca^2+^]_i_ signalization and intercellular connectivity. Finally, further studies are needed to understand the temporal evolution of the described changes during progression of diabetes, the mechanistic relationship between the observed changes in [Ca^2+^]_i_ signals and the other functional parameters, such as changes in electrical activity and insulin secretion (47, 48), as well as with pathophysiological factors characterizing beta cell dysfunction (49).

## Acknowledgements

We thank the Human Organ Procurement and Exchange (HOPE) program and Trillium Gift of Life Network (TGLN) for their work in procuring human donor pancreas for research. We also thank Drs. James Shapiro and Tatsuya Kin (University of Alberta Clinical Islet Program) for contributing some islet preparations for this study. Finally, we especially thank the organ donors and their families for their kind gift in support of diabetes research.

## Author Contributions

M.G., R.Y., H.L., P.E.M., and A.S. designed the study and contributed to conceptualization. R.Y. and H.L. performed the experiments. M.G. analyzed the data. M.G., R.Y., P.E.M., and A.S. contributed to the interpretation of the results. M.G. wrote the original draft of the manuscript and prepared the figures. P.E.M. and A.S. contributed to discussion and reviewed and edited the manuscript. P.E.M. and A.S. were responsible for supervision, funding acquisition, and project administration. A.S. is the guarantor of this work and, as such, had full access to all the data in the study and takes responsibility for the integrity of the data and the accuracy of the data analysis.

## Prior Presentation

Parts of this study were presented in abstract form at the Human Islet Research Network (HIRN) annual meetings, 28 April-1 May 2019 and 30 September-1 October 2020.

## Supplementary figures

**Supplementary figure S1.**
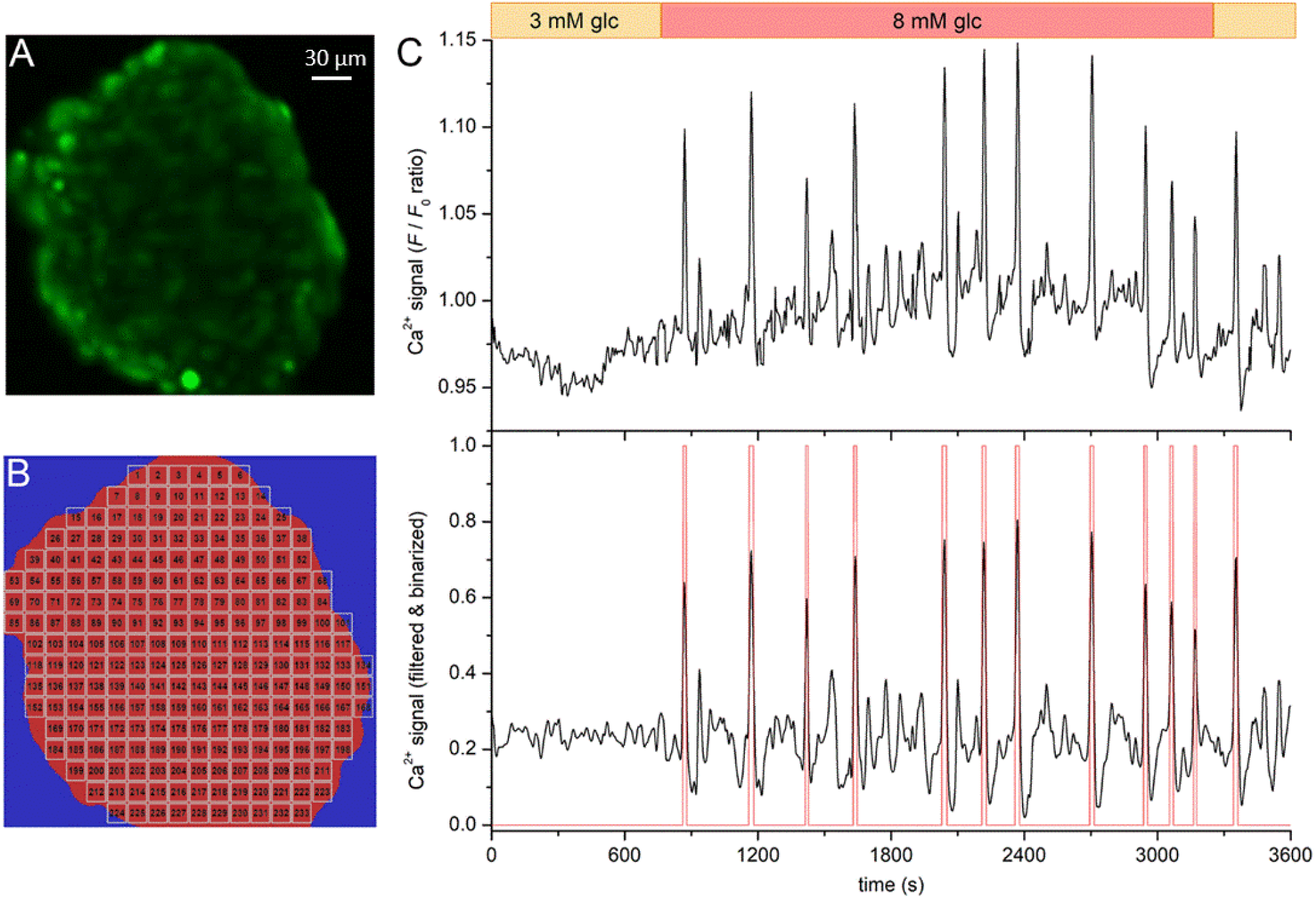
Methodology to extract [Ca^2+^]_i_ dynamics in isolated human islets. An image of an isolated islet loaded with Fluo-4 (A). Binarized image (red area on blue background) of the islet that is divided into square islet subregions (ISRs) with a linear size of 15 µm (white squares), from which the [Ca^2+^]_i_ signals were extracted (B). Unprocessed [Ca^2+^]_i_ signal from one ISR (upper row) and the corresponding high-pass filtered [Ca^2+^]_i_ trace (black line) along with the binarized activity (red line) (C). A cut-off frequency 0.02-0.04 Hz of the high-pass filter was used together with a binarization threshold that was determined based on the standard deviation of the filtered time series. After binarization, a smoothing-based refinement of the binarized signals was preformed to remove artefacts.

**Supplementary figure S2.**
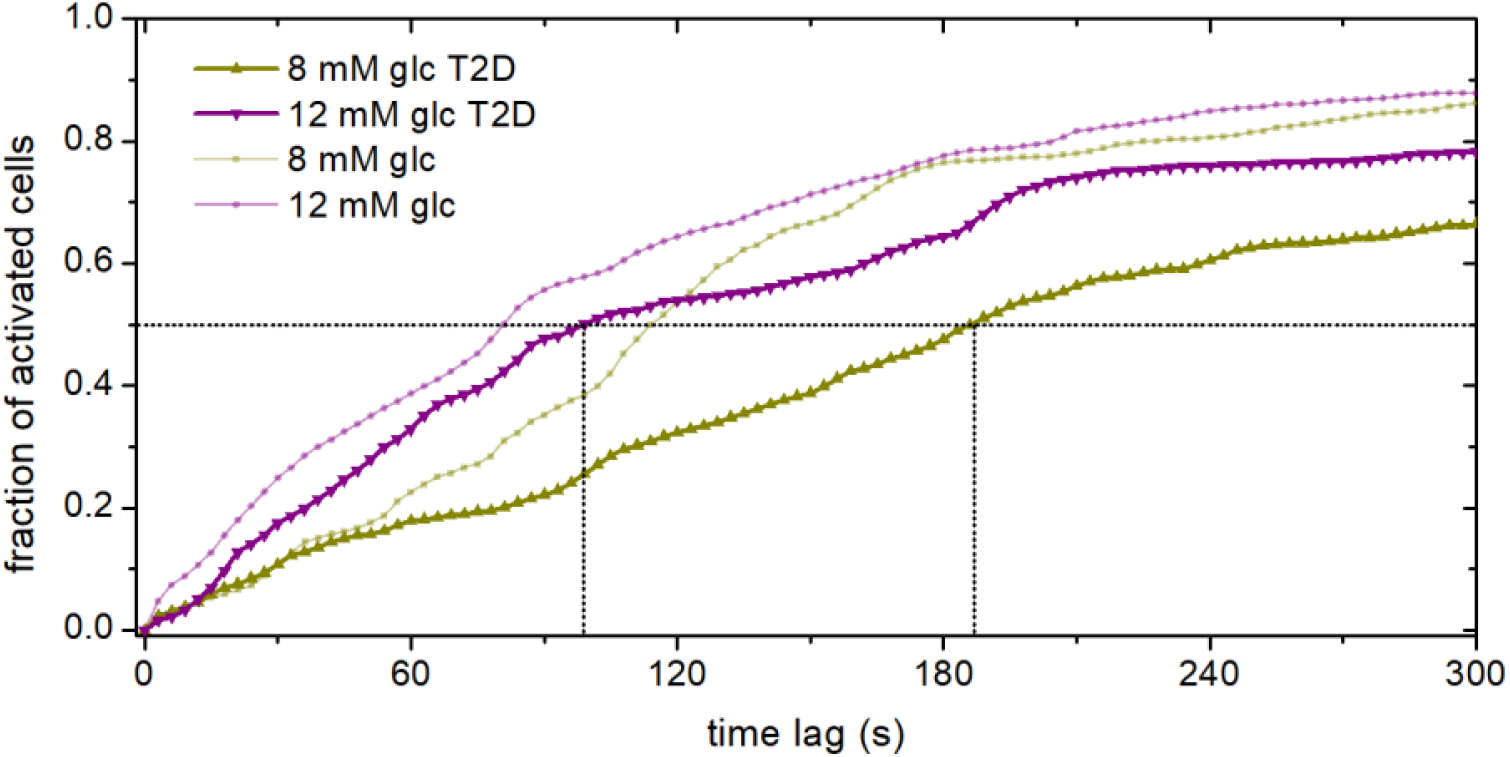
Activation is faster in higher glucose and delayed in islets from donors with type 2 diabetes. Cumulative activation of ISRs after the rise in glucose concentration from substimulatory 3 mM to stimulatory 8 mM or 12 mM for islets from control donors and donors with type 2 diabetes. An ISR was considered activated the first time a [Ca^2+^]_i_ oscillation occurred after rise in glucose stimulation. Both thick lines denote the average over all islets from donors with type 2 diabetes, whereas the thin light lines indicate the average behavior from non-diabetic donors, for easier comparison. Dotted lines indicate t_50_.

**Supplementary figure S3.**
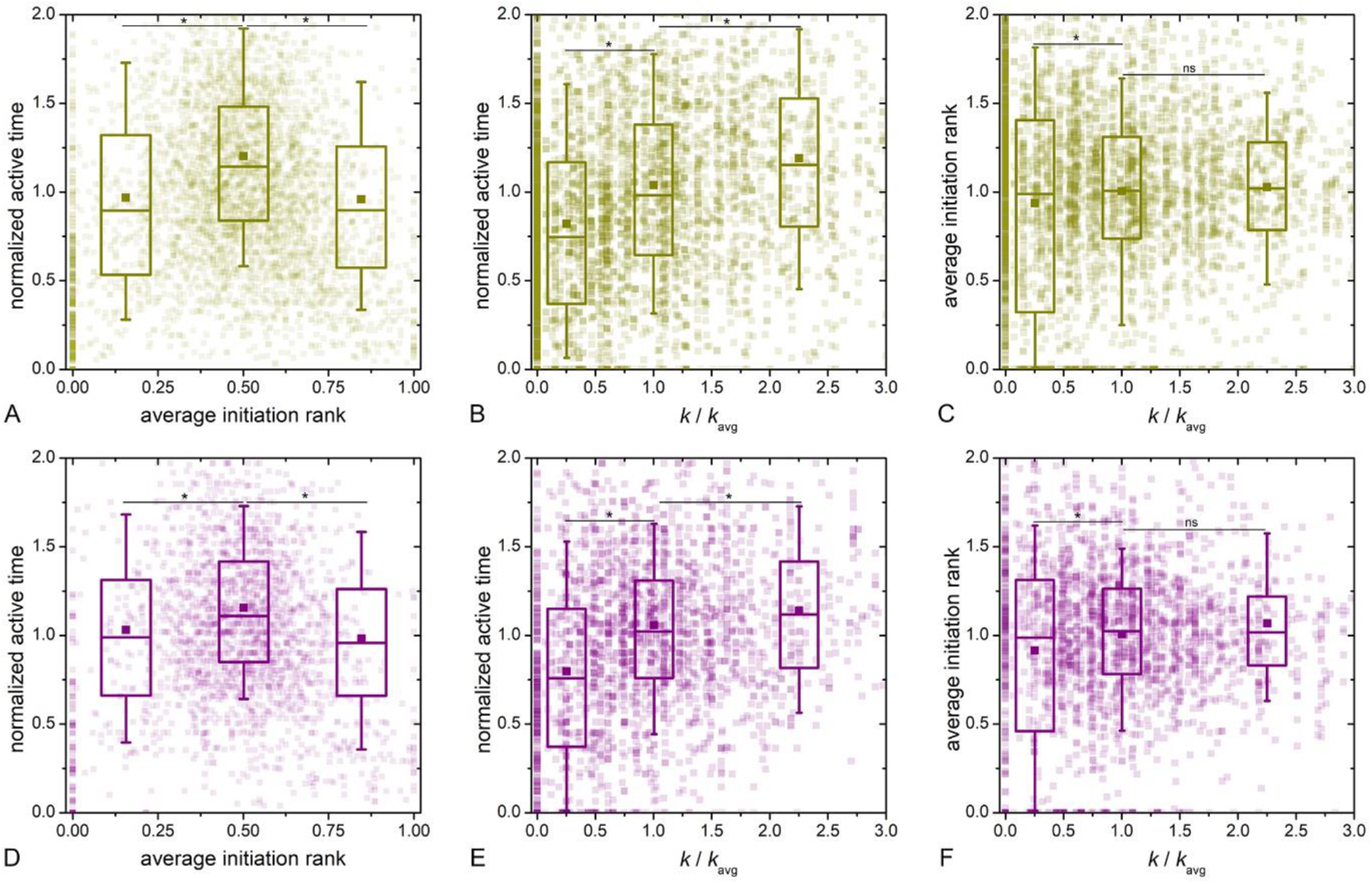
The relationships between active time, average initiation rank, and relative node degree for all islets from donors with type 2 diabetes. Transparent dots indicate individual ISRs, whereas the box-plots denote the lowest, the intermediate, and the highest one third of initiation rank parameter values (A and D) or the least connected cells (*k*/*k*_avg_≤0.5), intermediately connected cells (0.5< *k*/*k*_avg_ ≤1.5), and the most connected cells (*k*/*k*_avg_ >1.5) (B, C, E and F). Box charts are defined as in Figures 1-4. The results were obtained on the basis of 3971 and 2256 ISRs for 8 mM and 12 mM glucose, respectively.

**Supplementary table 1.** Basic characteristics, islet insulin secretion, and insulin content in donors with and without type 2 diabetes.

| Record ID | Age | Sex | Donation type | BMI | HbA1c | T2D | 1 mM glucose (pg/ml) |  |  | 10 mM glucose (pg/ml) |  |  | Content (ng/ml) |  |  |
| --- | --- | --- | --- | --- | --- | --- | --- | --- | --- | --- | --- | --- | --- | --- | --- |
| R234 | 50 | Female | NDD - Neurological | 31,7 | 5,7 | No | 482,703 | 883,181 | 320,667 | 4585,47 | 9378,6 | 4817,62 | 552,1768 | 704,6044 | 699,3864 |
| R235 | 53 | Female | NDD - Neurological | 24,5 | 5,7 | No | 1273,248 | 2129,952 | 809,231 | 5307,08 | 2712,37 | 5445,56 | 875,9488 | 1020,7028 | 2017,2964 |
| R272 | 56 | Male | NDD - Neurological | 26,9 | 5,6 | No | 2076 | 524 | 1987 | 3236 | 2870 | 3741 | 1459,028 | 1156,94 | 1555 |
| R280 | 17 | Female | NDD - Neurological | 26,3 | 5,2 | No | 364 | 614 | 394 | 4938 | 6868 | 5010 | 557,892 | 438,974 | 277,597 |
| H2169 | 26 | Male | NDD - Neurological | 22,1 | 5,2 | No | n.a. | n.a. | n.a. | n.a. | n.a. | n.a. | n.a. | n.a. | n.a. |
| H2171 | 54 | Male | NDD - Neurological | 33,4 | 6,3 | No | n.a. | n.a. | n.a. | n.a. | n.a. | n.a. | n.a. | n.a. | n.a. |
| R236 | 51 | Male | DCD - Cardio-Circulatory | 35,3 | 8,6 | Yes | 118,806 | 7,512 | 8,685 | 116,33 | 46,75 | 131,96 | 9,1672 | 3,0256 | 0,438 |
| R240 | 55 | Female | NDD - Neurological | 30,9 | 6,9 | Yes | 1472,052 | 1934,836 | 382,52 | 5586,53 | 5296,78 | 3022,2 | 1270,9792 | 1094,3584 | 665,596 |
| R244 | 48 | Female | NDD - Neurological | 30,4 | 7,5 | Yes | 573,07 | 818,691 | 1232,543 | 3102,5 | 2368,47 | 4706,34 | 568,4272 | 668,5328 | 1381,4516 |
| R262 | 54 | Female | NDD - Neurological | 22,4 | 5,6 | Yes | 318,084 | 120,184 | 196,558 | 1393,75 | 2341,94 | 3880,45 | 1538,0808 | 1538,0808 | 824,2572 |
| R263 | 71 | Male | NDD - Neurological | 38,3 | 7,6 | Yes | 1216,839 | 2357,935 | 1504,661 | 5654,84 | 6574,57 | 7752,23 | 1080,4516 | 2085,5264 | 1935,0264 |
| R307 | 43 | Male | NDD - Neurological | 25,4 | 6,5 | Yes | 550 | 631 | 561 | 1812 | 2067 | 24633 | 567,792 | 1039,486 | 548,566 |

**Supplementary video S1**. Recording of a human islet and the corresponding animation of binarized spatiotemporal [Ca^2+^]_i_ activity in a representative islet stimulated with the 3-12-3 mM glucose protocol.

